# The physiological level of rNMPs present in mtDNA does not compromise its stability

**DOI:** 10.1101/746719

**Authors:** Paulina H. Wanrooij, Phong Tran, Liam J. Thompson, Sushma Sharma, Katrin Kreisel, Clara Navarrete, Anna-Lena Feldberg, Danielle L. Watt, Anna Karin Nilsson, Martin K. M. Engqvist, Anders R. Clausen, Andrei Chabes

## Abstract

Ribonucleotides (rNMPs) incorporated in the nuclear genome are a well-established threat to genome stability and can result in DNA strand breaks when not removed in a timely manner. However, the presence of a certain level of rNMPs is tolerated in mitochondrial DNA (mtDNA), although aberrant mtDNA rNMP content has been identified in disease models. We investigated the effect of incorporated rNMPs on mtDNA stability over the mouse lifespan and found that the mtDNA rNMP content increased during early life. The rNMP content of mtDNA varied greatly across different tissues and was defined by the rNTP/dNTP ratio of the tissue. Accordingly, mtDNA rNMPs were nearly absent in *SAMHD1*^*−/−*^ mice that have increased dNTP pools. The near absence of rNMPs did not, however, appreciably affect mtDNA copy number or the levels of mtDNA molecules with deletions or strand breaks in aged animals near the end of their lifespan. The physiological rNMP load therefore does not contribute to the progressive loss of mtDNA quality that occurs as mice age.

## Introduction

The mitochondrial genome in mammalian cells is a circular 16 kb double-stranded DNA molecule present in multiple copies per cell. This mitochondrial (mt)DNA encodes key subunits of the respiratory chain complexes that synthesize the majority of the cell’s ATP. Consequently, partial loss of the number of copies of mtDNA from the cell (i.e. depletion) or defects in the quality of the DNA can affect energy production and manifest as human disease^1^. MtDNA quality also decreases with age due to the progressive accumulation of deletions and point mutations^2–5^, which might contribute to the physiology of aging^6^.

Ribonucleotides (rNMPs) incorporated into DNA are an important threat to genome stability. Although replicative DNA polymerases are highly selective for dNTPs, the large excess of rNTPs over dNTPs in the cell results in the incorporation of some rNMPs during DNA replication ^7,8^. Due to their reactive 2′-hydroxyl group, these rNMPs increase the risk of strand breaks by several orders of magnitude^9^. Furthermore, the presence of rNMPs in DNA can alter its local structure and elasticity^10–12^, thus interfering with processes such as replication, transcription, or DNA repair. To prevent these negative effects of incorporated rNMPs on genome stability, they are removed from nuclear DNA (nDNA) by the ribonucleotide excision repair pathway, which is initiated by cleavage at the incorporated rNMP by RNase H2^8,13^. RNase H2 is essential for genome stability in mammals, and its absence results in increased single-stranded DNA (ssDNA) breaks and activation of the DNA damage response^14,15^. However, incorporated rNMPs can also have positive roles in the maintenance of genome stability, at least when present transiently. For example, the unremoved rNMPs in the nascent leading strand have been proposed to act as a mark of the nascent strand during mismatch repair^16,17^, and rNMPs incorporated at double-strand DNA breaks might promote break repair^18^.

In contrast to nDNA, mtDNA contains persistent rNMPs that are not removed after incorporation by the replicative mtDNA polymerase Pol ɣ (reviewed in ^19^). Human Pol ɣ is highly selective against using rNTPs as substrates and thus incorporates fewer rNMPs than nuclear replicative DNA polymerases do^20–22^. Once incorporated, however, the single rNMPs persist in mtDNA because mitochondria lack repair pathways for their efficient removal^23,24^. Although it is clear that a certain level of rNMPs is tolerated in mtDNA, their effect on mtDNA stability is not well understood. Interestingly, the absolute and relative rNMP content of mtDNA is altered in cell lines from patients with mtDNA depletion syndrome (MDS)^24^. Furthermore, aberrant rNMP incorporation has been proposed to play a role in the mtDNA deletions and depletion observed in mice lacking the mitochondrial inner membrane protein MPV17^25^.

We report here that the mtDNA rNMP content varies considerably between different mammalian tissues and is determined by the rNTP/dNTP ratio in the respective tissues. In light of the above reports connecting aberrant rNMP content to mtDNA instability^24,25^ and, on the other hand, the notion that incorporated rNMPs might play a positive role in genome maintenance, we asked whether the normal, physiological level of rNMPs affects mtDNA stability over the lifespan of the mouse. To this end, we used a mouse model that is deficient in the dNTP triphospohydrolase (dNTPase) SAMHD1 and consequently has higher levels of dNTPs than wild-type (wt) mice. In accordance with the decreased rNTP/dNTP ratio in *SAMHD1*^*−/−*^ animals, we observed a drastic reduction in mtDNA rNMP content. This change, however, did not affect the occurrence of mtDNA deletions during aging nor did it alter the mtDNA copy number or the number of strand breaks in aged animals. These results demonstrate that normal levels of rNMPs have no apparent detrimental or beneficial effects on mtDNA integrity over the lifespan of the mouse.

## Results

### The rNMP content of mtDNA varies between tissues

We examined the rNMP content of mtDNA isolated from mouse tissues by comparing the electrophoretic mobility of untreated and alkali-treated DNA on a denaturing gel. To validate that the extent of alkali sensitivity provides an accurate readout of the rNMP content of mtDNA, we treated samples from mouse liver, heart, and skeletal muscle (gastrocnemius (GAS)) either with alkali or with recombinant RNase HII, an enzyme that cleaves the DNA backbone at incorporated rNMPs, and analyzed the resulting mtDNA using a probe against the *COX1* gene under denaturing conditions that cause separation of the two DNA strands. Both treatments caused a similar decrease in the median length of mtDNA fragments (Fig. 1a, b), confirming that mtDNA from these three tissues contains rNMPs and that most alkali-sensitive sites are rNMPs rather than abasic sites, which are also sensitive to KOH. Thus, we used alkali treatment in further experiments as an assay for rNMPs in mtDNA.

**Figure 1.**
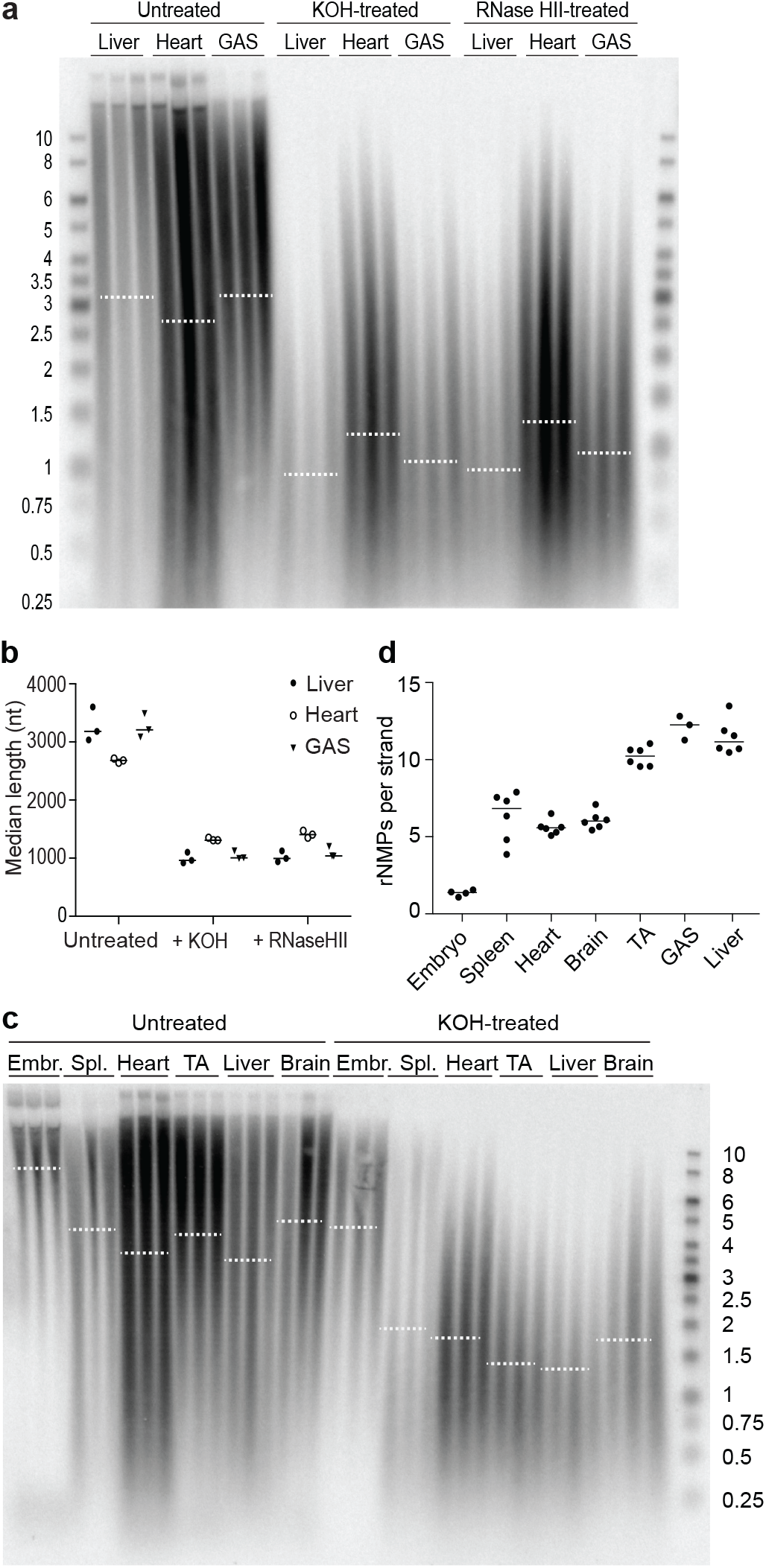
The rNMP content of mtDNA varies across tissues. **a)** DNA isolated from liver, heart, and the gastrocnemius muscle (GAS) was separated by electrophoresis on a denaturing gel either before (Untreated) or after treatment with alkali (KOH) or RNase HII, both of which cleave the DNA backbone at incorporated rNMPs. MtDNA was visualized using a *COX1* probe. Each sample corresponds to an individual mouse. Dotted lines represent the median length of the mtDNA in each group. **b)** The median length of the mtDNA before (Untreated) or after treatment with alkali (+ KOH) or RNase HII was quantified from the distribution and intensity of the signal in Fig. 1a. Filled circles: liver; empty circles: heart; triangles: GAS. Each symbol represents an individual mouse, and lines indicate the median. **c)** Untreated and KOH-treated DNA from various adult mouse tissues and from embryos was separated on a denaturing gel, and mtDNA was visualized by Southern blotting as above. Each sample lane represents an individual mouse. Three independent experiments were performed, and a representative image is shown. Dotted lines represent the median. See Fig. S1a–c for the same blot with other probes. **d)** The length difference between untreated and alkali-treated samples from three separate experiments including the one shown in Fig. 1c was used to compute the number of rNMPs per single strand of mtDNA in embryos and in various adult tissues. Each dot represents an individual mouse, and lines indicate the median. The sizes of DNA ladder bands are indicated in kb. See also Fig. S1.

Next, we compared the mtDNA rNMP content in embryos and across various tissues from adult mice. Alkaline hydrolysis resulted in substantially shorter mtDNA fragments than in the untreated samples from liver and skeletal muscle (tibialis anterior (TA) and GAS), whereas the mtDNA from embryos was much less affected (Fig. 1c). By comparing the median lengths of the DNA in the untreated and alkali-treated samples, we estimated the number of rNMPs incorporated per strand of mtDNA. This revealed clear differences in mtDNA rNMP content between tissues (Fig. 1c, d). Spleen, heart, and brain mtDNA contained ~6 rNMPs per strand of mtDNA, whereas liver and both the TA and GAS muscles of the hind leg contained 10–12 rNMPs per strand. MtDNA from embryos had the lowest rNMP content, with only ~1 rNMP per strand of mtDNA. Stripping and re-hybridization of the *COX1* blot in Fig. 1c with a probe annealing to another region of the mtDNA – the D-loop, a triple-stranded region formed by stable incorporation of a short DNA strand known as 7S DNA – indicated that both regions have similar rNMP content (Fig. S1a; quantification was not carried out due to the interfering signal of the 7S DNA). Finally, Southern blot analysis using strand-specific ssDNA probes revealed no apparent strand bias in rNMP content between the H- and L-strand of mtDNA (Fig. S1b). As expected, nDNA from the various tissues was not affected by alkaline hydrolysis (Fig. S1c).

### MtDNA rNMP content correlates with the rNTP/dNTP ratio

While some studies have found that the ratio of free rNTPs to dNTPs determines the frequency of rNMP incorporation in mtDNA of the yeast *Saccharomyces cerevisiae*^23^ and of cultured human cells^24^, others have reported changes in the mtDNA rNMP content that did not correspond to changes in rNTP/dNTP ratios^25^. We therefore sought to determine whether the differences in mtDNA rNMP content we observed in various tissues correlated with differing rNTP/dNTP ratios. We measured the levels of individual rNTPs and dNTPs in mouse embryos and in spleen and skeletal muscle from adult animals by high-performance liquid chromatography and calculated the ratio of each rNTP to its corresponding dNTP (Table 1). Of the four rNTP/dNTP pairs, the rATP/dATP ratio was highest, in accordance with the abundance of rATP in most cell types^7,26^, and it varied the most between tissue types – embryonic tissue contained a 470-fold excess of rATP over dATP, in spleen there was an 800-fold excess, and in skeletal muscle there was a 37,000-fold excess. This high ratio observed in skeletal muscle exceeds the ability of the mitochondrial DNA polymerase, Pol ɣ, to efficiently distinguish rATP from dATP (with a discrimination factor of 9,300-fold^21^). The ratio of the other three rNTP/dNTP pairs varied considerably less between tissues and remained well below the value of the Pol ɣ discrimination factor for each pair^21^. The rATP/dATP ratio might therefore have the greatest impact on the rNMP content of mtDNA. Indeed, comparison of embryos, spleen, and skeletal muscle confirmed that the rNMP content of mtDNA reflects the magnitude of the rATP/dATP ratio (Fig. 1c, Table 1). In essence, the mtDNA rNMP content is therefore a readout of the rNTP/dNTP ratio of each tissue.

**Table 1.**
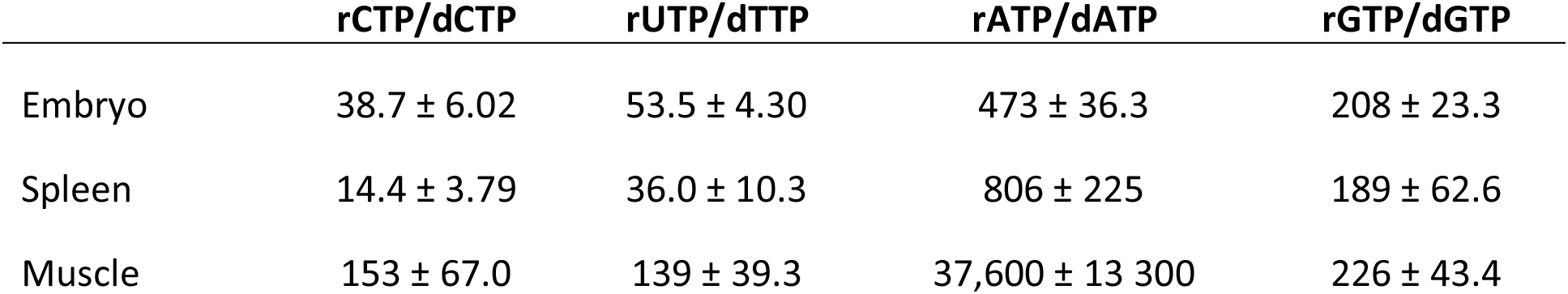
rNTP/dNTP ratios from mouse embryos and adult spleen and skeletal muscle. The means and standard deviations of three or four independent samples are shown.

|  | rCTP/dCTP | rUTP/dTTP | rATP/dATP | rGTP/dGTP |
| --- | --- | --- | --- | --- |
| Embryo | 38.7 ± 6.02 | 53.5 ± 4.30 | 473 ± 36.3 | 208 ± 23.3 |
| Spleen | 14.4 ± 3.79 | 36.0 ± 10.3 | 806 ± 225 | 189 ± 62.6 |
| Muscle | 153 ± 67.0 | 139 ± 39.3 | 37,600 ± 13 300 | 226 ± 43.4 |

### The rNMP content of mtDNA increases during development

Based on Fig. 1c, the mtDNA from embryos contained very few incorporated rNMPs (~1 rNMP per strand of mtDNA), while the number of rNMPs in all analyzed adult tissues was clearly higher. This suggests temporal changes in mtDNA rNMP content during development. To follow changes in rNMP incorporation at various ages, we analyzed the mtDNA from embryos and from the skeletal muscle of pups (15 days old), adults (13 weeks old), and aged (2 years old) animals. As previously observed, embryonic mtDNA contained very few rNMPs (Fig. 2a, b; see Fig. S2a for nDNA). Furthermore, mtDNA rNMP content increased from ~5 per strand in the skeletal muscle of pups to ~8.5 per strand in adult animals. No difference was observed, however, between the number of rNMPs in adult and aged mice, demonstrating that rNMP content in skeletal muscle mtDNA does not increase during aging. Similar to the observations in skeletal muscle, the rNMP content of mtDNA from the heart was also higher in adult mice (~5 rNMPs per strand) than in pups (~1 rNMP per strand), but in contrast to the skeletal muscle there was little difference between embryos and pups (Fig. 2c, Fig. S2b-c). These results demonstrate that the mtDNA rNMP content increases during early life but not during aging, and that there are temporal differences in rNMP accumulation between skeletal and heart muscle (compare Fig. 2b and Fig. 2c).

**Figure 2.**
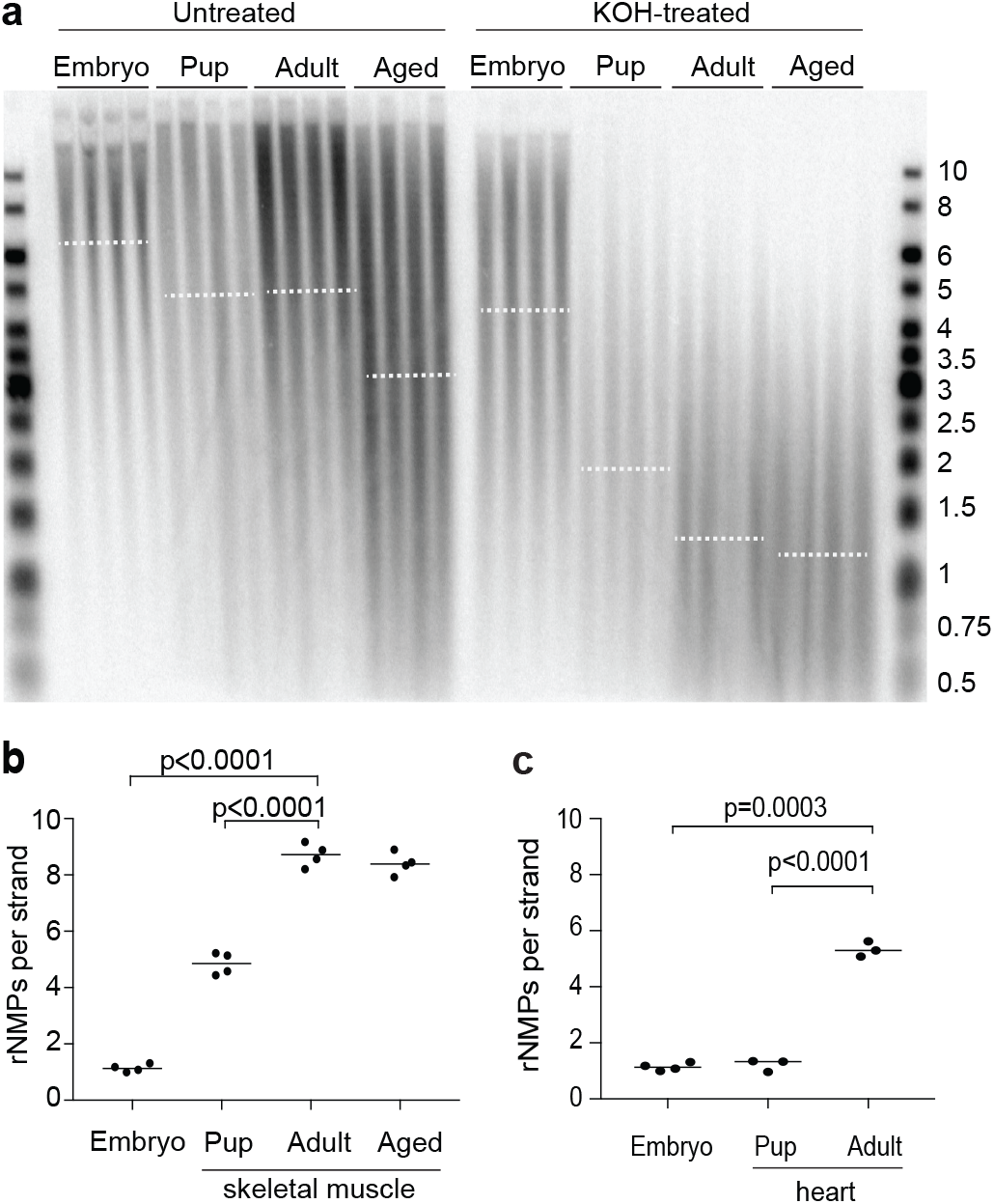
The rNMP content of mtDNA increases during development. **a)** DNA from embryos and from the TA muscle of pups, adult, and aged animals was left untreated or hydrolyzed with alkali (KOH-treated). Samples were separated on a denaturing gel, and mtDNA was visualized using a *COX1* probe. Each sample corresponds to an individual mouse. The experiment was performed twice, and a representative blot is shown. Dotted lines represent the median. **b)** The length difference between untreated and alkali-treated samples in Fig. 2a was used to compute the number of rNMPs per single strand of mtDNA. The p-values of statistically significant deviations from the adult are indicated (Welch’s t-test; n = 4), and the line indicates the median. **c)** The length difference between untreated and KOH-treated heart DNA shown in Fig. S2b was used to compute the number of rNMPs per single strand of mtDNA. Note that the embryo values are the same as in Fig. 2b. The p-values of statistically significant differences between adult heart rNMP content and the other samples are indicated (Welch’s t-test; n = 3), and the horizontal lines indicate the median for each group. The sizes of the bands in the DNA ladder are indicated in kb. See also Fig. S2.

### Mice with elevated dNTP pools have fewer rNMPs in their mtDNA

We and others have previously demonstrated that mice deficient in the dNTP triphosphohydrolase SAMHD1 have elevated dNTP levels^27–29^. Because the mtDNA rNMP content correlates directly with the rNTP/dNTP ratio, we reasoned that mtDNA from *SAMHD1*^*−/−*^ mice with increased dNTP levels should contain fewer rNMPs. To test our hypothesis, we first examined the rNMP content of skeletal muscle mtDNA from wt and *SAMHD1*^*−/−*^ mice. The number of rNMPs in mtDNA from GAS and TA muscle of *SAMHD1*^*−/−*^ animals (fewer than 1 rNMP per strand) was much lower than that of wt (~9 and ~8 rNMPs per strand) (Fig. 3a, b). In contrast, the mtDNA rNMP content of muscle from *SAMHD1*^*+/−*^ heterozygotes was not significantly affected, which was in accordance with the mild effect of heterozygosity on their rNTP/dNTP ratios^28^. SAMHD1-deficiency had no effect on the sensitivity of nDNA to alkaline hydrolysis (Fig. S3a). An equally striking decrease in rNMP content was observed in the mtDNA from *SAMHD1*^*−/−*^ liver, and this persisted in old adult mice (1 year old; Fig. 3c, d; see Fig. S3b for nDNA blot). We also examined heart mtDNA from old adults and found fewer rNMPs in *SAMHD1*^*−/−*^ mice than in wt mice (Fig. S3c–e), confirming the effect even in a tissue with a lower rNMP content than that of skeletal muscle and liver.

**Figure 3.**
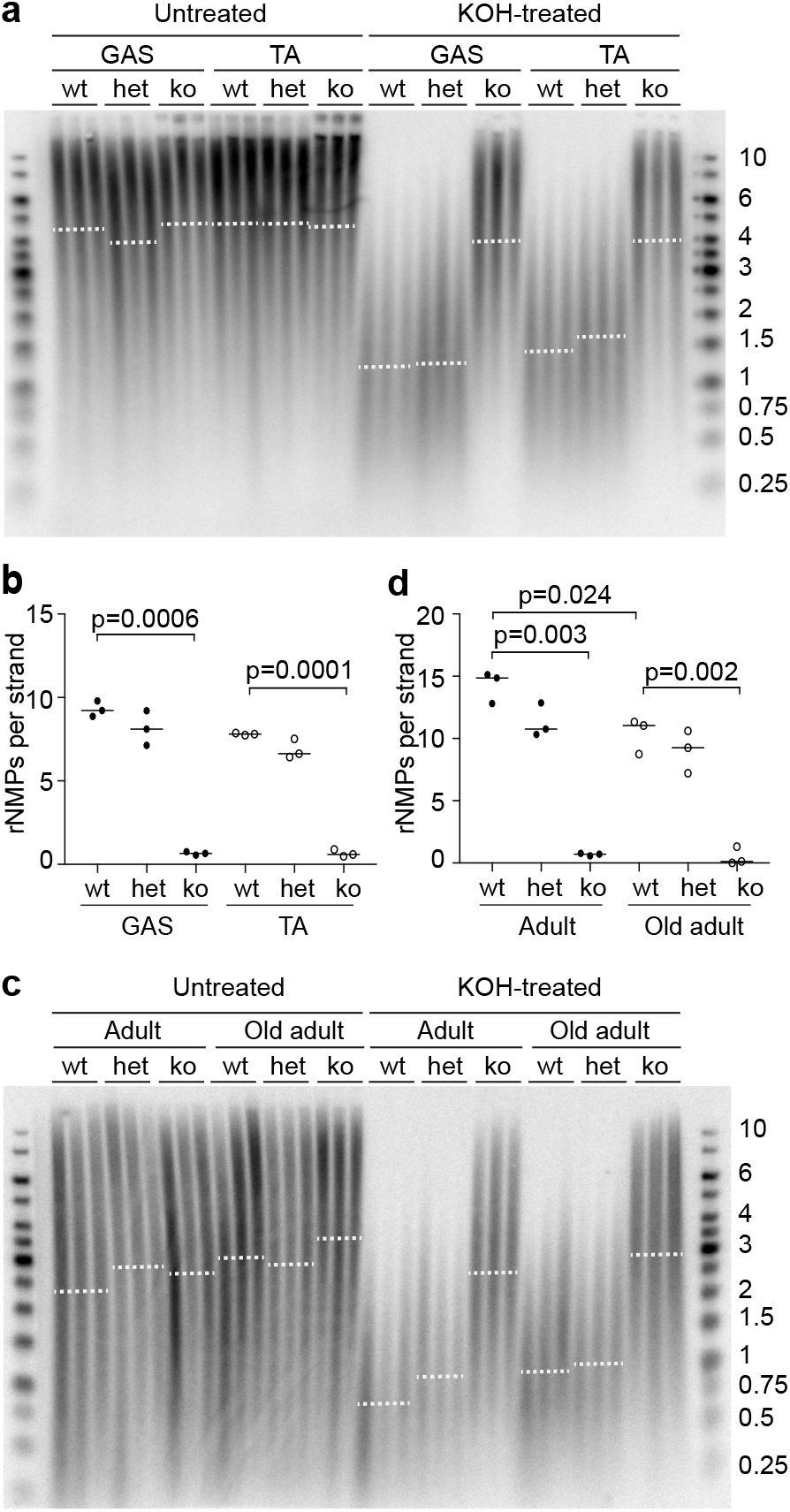
MtDNA from *SAMHD1*^*−/−*^ mice contains very little rNMPs. **a)** Untreated and alkali-treated DNA from skeletal muscle of adult wt and SAMHD1-deficient mice was analyzed on a denaturing gel, and mtDNA was visualized using a *COX1* probe. Each sample corresponds to an individual mouse. The experiment was carried out twice, and a representative image is shown. GAS, gastrocnemius; TA, tibialis anterior; wt, wild type; het, *SAMHD1*^*+/−*^; ko, *SAMHD1*^*−/−*^. Dotted lines represent the median. **b)** The length difference between untreated and alkali-treated mtDNA in Fig. 3a was used to compute the content of rNMPs per single strand of mtDNA. The p-values of statistically significant deviations from the wt are indicated (Welch’s t-test; n = 3), and the horizontal lines indicate the median for each group. **c)** Untreated or KOH-treated DNA from the livers of 13-week-old (adult) or 1-year-old (old adult) wt and SAMHD1-deficient mice was analyzed on a denaturing gel, and mtDNA was visualized as above. Each sample lane corresponds to an individual mouse. The experiment was performed three times, and a representative experiment is shown. wt, wild type; het, *SAMHD1*^*+/−*^; ko, *SAMHD1*^*−/−*^. Dotted lines represent the median. **d)** The length difference between untreated and alkali-treated mtDNA shown in Fig. 3c was used to compute the number of rNMPs per single strand of mtDNA. The p-values of statistically significant differences from the wt adult group are indicated (Welch’s t-test; n = 3), and horizontal lines indicate the median for each group. The sizes of bands in the DNA ladder are indicated in kb. See also Fig. S3.

To identify the rNMPs incorporated into the mtDNA of wt and *SAMHD1*^*−/−*^ animals, we used the HydEn–seq method^30^. In this method, KOH-treated samples are sequenced to uncover the identity and relative proportions of the four individual rNMPs. Consistent with the high rATP/dATP ratio in most tissues (Table 1; ^7,26^) and with previous reports on mtDNA from solid mouse tissues^25^, the most commonly occurring rNMP in the mtDNA from the liver of wt animals was rAMP, which comprised ~60% of all rNMPs (Fig. 4a). In comparison, rGMP comprised ~20%, and rUMP and rCMP each comprised ~10% of the total incorporated rNMPs. In contrast, the relative proportion of each individual dNMP at random background nicks, as determined by sequencing of 5′-ends in untreated (KCl-treated) samples, was close to 25% (Fig. S4a). In *SAMHD1*^*−/−*^ animals, mtDNA contains very few rNMPs (Fig. 3). Accordingly, the proportions of the four individual bases determined by HydEn-seq in the mtDNA of *SAMHD1*^*−/−*^ animals approached 25% (Fig. 4a), similar to the proportions of dNMPs at random background nicks (Fig. S4a). The similar proportion of bases at alkali-induced ends and at 5′-ends suggests that the majority of the “rNMPs” in the mtDNA of *SAMHD1*^*−/−*^ animals are in fact random background nicks rather than actual rNMPs. In contrast to the mtDNA, deletion of SAMHD1 did not alter the identity of the rNMPs in nDNA (Fig. 4b). Like the mtDNA in *SAMHD1*^*−/−*^ animals, the nDNA of wt and *SAMHD1*^*−/−*^ mice contained virtually no rNMPs (Fig. S3b). Therefore, the data obtained for nDNA similarly represent a low level of 5′-ends due to random background nicks. Consistent with this interpretation, the proportion of each “rNMP” in nDNA was ~25%, comparable to the proportion of each dNMP at 5′-ends as determined by 5′-end sequencing (Fig. S4b). Finally, we analyzed the rNMPs incorporated into each individual strand of the mtDNA. No strand bias was found in the incorporation of rNMPs into mtDNA from wt or *SAMHD1*^*−/−*^ when analyzed by HydEn–seq (Fig. 4c), which was in accordance with the Southern blot analysis (Fig. S1b). Similar analysis of nDNA showed no difference between the two strands (Fig. S4c). Taken together, these data suggest that the dNTP pool alterations observed in *SAMHD1*^*−/−*^ animals reduce both the amount of rNMPs incorporated into the mitochondrial genome and the relative proportion of rAMP that is incorporated.

**Figure 4.**
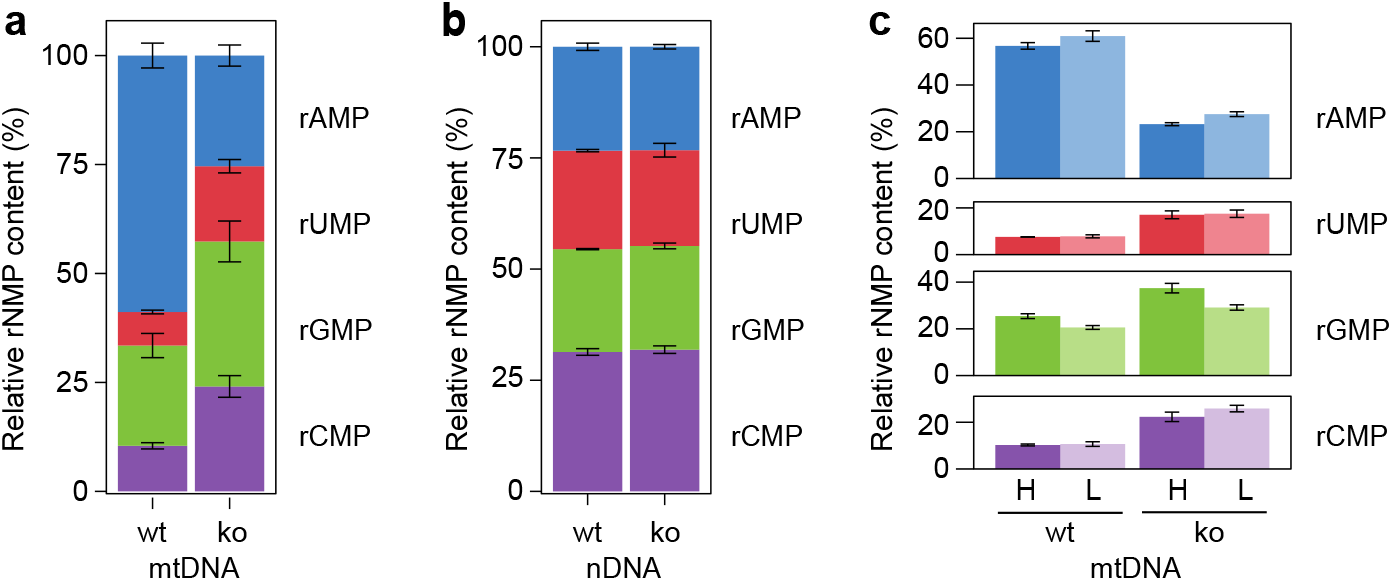
The proportions of different rNMPs in mtDNA is altered in *SAMHD1*^*−/−*^ animals. The base identities of rNMPs incorporated in the mitochondrial and nuclear genomes from the liver of adult wt and *SAMHD1*^*−/−*^ (ko) mice were analyzed by HydEn-Seq. Three wt and four *SAMHD1*^*−/−*^ mice were analyzed. **a)** The proportion of each rNMP (rAMP, rUMP, rGMP, and rCMP) as a percentage of the total rNMPs present in the mtDNA was plotted for wt and *SAMHD1*^*−/−*^ mice. Error bars represent the standard deviation. **b)** The proportion of each rNMP as a percentage of the total rNMPs present in the nDNA was plotted for wt and *SAMHD1*^*−/−*^ mice. Error bars represent the standard deviation. **c)** The proportion (%) of each rNMP in each mtDNA strand (H, L) in wt and *SAMHD1*^*−/−*^ mice. Error bars represent the standard deviation. See also Fig. S4.

### Reduction of rNMPs does not affect mtDNA integrity in aged skeletal muscle

The presence of incorporated rNMPs in DNA is a serious threat to genome stability, as exemplified by the increased ssDNA breaks observed in nDNA in the absence of functional ribonucleotide excision repair^14,15^. On the other hand, rNMPs might play a functional role in DNA metabolism and thereby be present in mtDNA for a reason. To address the impact of rNMPs on the mitochondrial genome, we asked whether the reduction of rNMP content from physiological levels affects mtDNA stability over the course of the mouse lifespan. We were especially interested in studying the effects of a reduced rNMP load on the mtDNA of aged animals because of the reported age-dependent reduction in mtDNA integrity that might contribute to the physiology of aging^2,3,5,6^.

First, we determined the effect of *SAMHD1* deletion on mtDNA copy number by quantitative real-time PCR. The mtDNA copy number in adult *SAMHD1*^*−/−*^ mice was slightly higher than in wt, whereas no difference was observed between old adult or aged *SAMHD1*^*−/−*^ and wt animals (Fig. 5a). In liver, both adult and old adult *SAMHD1*^*−/−*^ animals had a higher mtDNA copy number compared to wt animals of the same age, while aged animals remained unaffected (Fig. S5a). Because increased dNTP pools are known to increase mtDNA copy number in yeast^23,31^, our primary interpretation of these data is that dNTP pools might limit mtDNA copy number in the muscle and livers of adult but not aged animals. Overall, copy number analysis revealed no striking effect of the reduction of rNMPs on mtDNA levels in aged animals that are close to the end of their lifespan.

**Figure 5.**
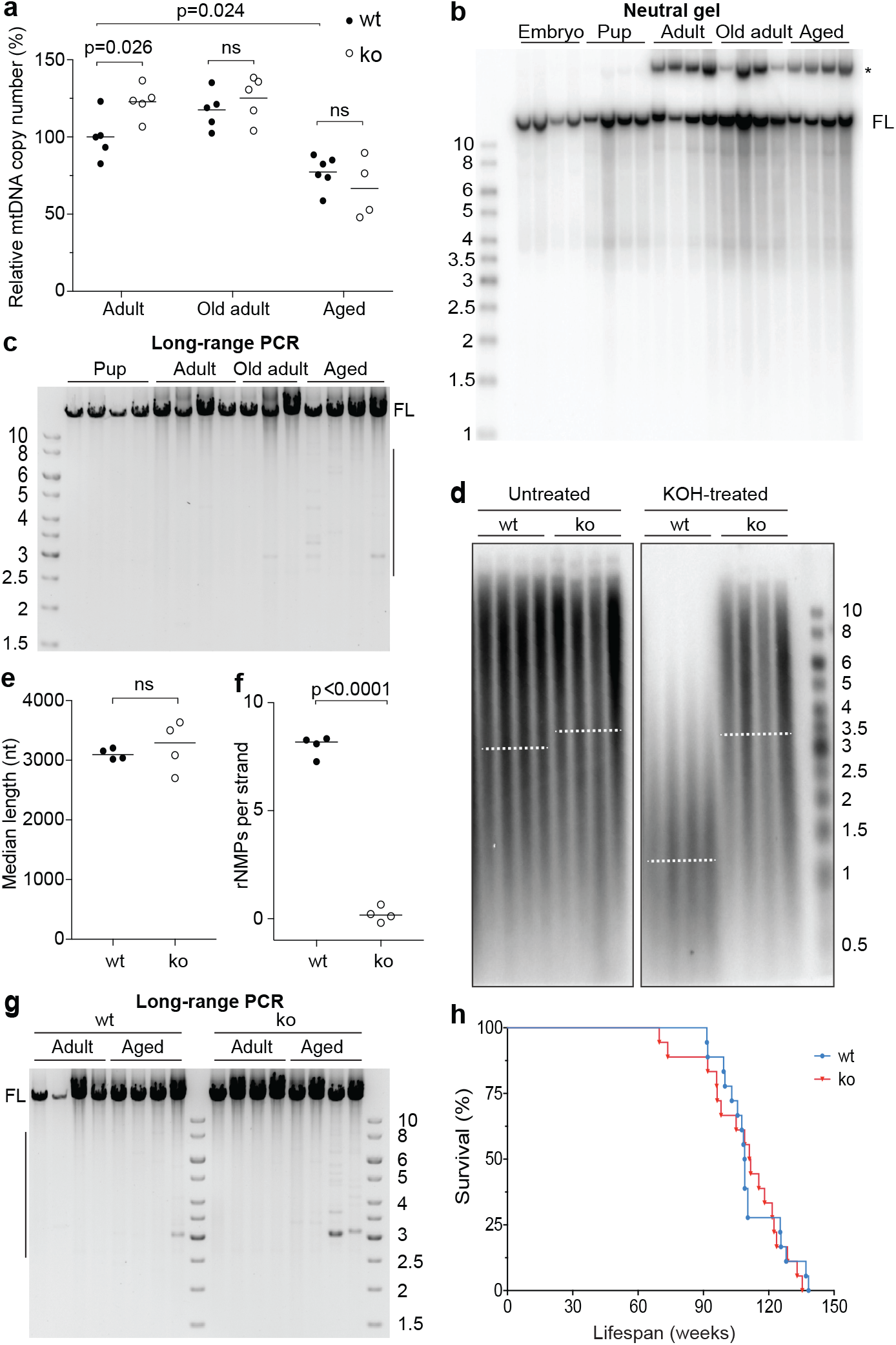
Deletion of *SAMHD1* does not affect mtDNA stability. **a)** MtDNA copy number in the TA muscle of 5 or 6 wt (filled dots) and *SAMHD1*^*−/−*^ (open dots) 13-week-old (adult), 1-year-old (old adult), and 2-year-old (aged) animals was determined by qPCR and normalized to the value for adult wt mice. The mean for each group is indicated by a horizontal line. The p-values were calculated using Welch’s t-test; ns, non-significant. **b)** DNA isolated from embryos and from the TA muscle of pups, adults, 1-year-old (old) adults, or aged animals was linearized with SacI endonuclease and separated on a neutral gel. MtDNA was visualized as above. Full-length mtDNA is indicated (FL); the asterisk denotes a higher-migrating species resistant to cleavage. **c)** Long-range PCR to detect deletions in mtDNA from the TA muscle of wt mice of various ages. Full-length product is indicated (FL). Only minor species containing deletions are observed in the mtDNA from old adults and aged animals, as indicated by the vertical line on the right-hand side of the gel. **d)** Untreated or alkali-treated DNA from skeletal muscle of aged wt and *SAMHD1*^*−/−*^ (ko) mice was analyzed on a denaturing gel, and mtDNA was visualized using a *COX1* probe. Each sample lane corresponds to an individual mouse, and dotted lines represent the median. **e)** The median length of the untreated mtDNA in samples from Fig. 5d is indicated by a horizontal line. The two groups were compared using Welch’s t-test (ns, non-significant; n = 4). **f)** The length difference between untreated and alkali-treated mtDNAs shown in Fig. 5d was used to compute the number of rNMPs per single strand of mtDNA. The horizontal lines indicate the median. The p-value of the statistically significant difference between the two groups was calculated by Welch’s t-test; n = 4. **g)** Long-range PCR was performed on mtDNA isolated from the TA muscle of adult and aged wt or *SAMHD1*^*−/−*^ (ko) mice. FL, full-length product; the vertical line indicates the size range of mtDNA molecules with deletions. **h)** Kaplan–Meier survival curve for wt and *SAMHD1*^*−/−*^ (ko) mice. Comparison of the curves by the log-rank (Mantel–Cox) test confirmed no statistically significant difference between the genotypes. The sizes of the bands in the DNA ladder are indicated in kb. See also Fig. S5.

Next, we assessed mtDNA for the presence of large deletions because alterations in mtDNA rNMP content have been reported to correlate with an increase in mtDNA deletions in old animals^25^. We first ensured that we could detect mtDNA deletions in wt animals. Although no mtDNA species corresponding to shorter than full-length molecules were detected in skeletal muscle from various ages of mice by Southern blot (Fig. 5b), a more sensitive long-range PCR assay that preferentially amplifies mtDNA molecules containing deletions revealed some deletion-containing species in aged animal muscle (Fig. 5c). This is in agreement with previous findings that mtDNA deletions accumulate during aging^2,4^. However, even though the rNMP content of *SAMHD1*^*−/−*^ skeletal muscle mtDNA remained very low even in aged animals (Fig. 5d-f; the control for nDNA is shown in Fig. S5b), wt and *SAMHD1*^*−/−*^ muscle had similar amounts of mtDNA molecules containing deletions (Fig. 5g). Furthermore, the comparable migration of untreated wt and *SAMHD1*^*−/−*^ mtDNA on the denaturing gel in Fig. 5d indicates that the frequency of ssDNA breaks was not appreciably altered by the low rNMP content of *SAMHD1*^*−/−*^ muscle (quantified in Fig. 5e). Finally, we observed no difference between the lifespans of *SAMHD1*^*−/−*^ and wt mice (Fig. 5g). Taken together, these results suggest that the almost complete absence of rNMPs in *SAMHD1*^*−/−*^ mtDNA did not have a significant impact on animal wellbeing or on mtDNA stability in aged animals as judged by mtDNA copy number, frequency of deletions, or ssDNA breaks. We conclude that the physiological level of rNMPs is neither noticeably harmful nor essential for mtDNA maintenance in mouse muscle tissue.

## Discussion

MtDNA has previously been reported to contain a relatively large number of rNMPs^32–34^, which, as we demonstrate here, vary in frequency between tissues and increase in number during early life. It is well established that failure to remove rNMPs from the mammalian nuclear genome results in frequent ssDNA breaks, increased replication stress, and activation of the p53-dependent DNA damage response that leads to death during embryonic development^14,15^. In contrast, the cell tolerates the presence of a significant number of rNMPs in mtDNA^20,24^. This tolerance of rNMPs in mtDNA might be due to the multicopy nature of mtDNA, its slower rate of replication, and the reverse transcriptase activity of Pol ɣ that allows it to bypass single rNMPs in the DNA template more effectively than nuclear replicative polymerases do^20,21^. Although a certain level of rNMPs in mtDNA is well tolerated, we hypothesized that their continuous presence over the course of the lifespan might have a negative impact on mtDNA integrity through decreased bypass by Pol ɣ^20,21^ and increased ssDNA breaks and thereby contribute to the loss of mtDNA integrity that occurs with age.

To investigate whether rNMPs contribute to the mtDNA deletions observed in aged mice, we turned to a mouse model with increased dNTP levels. We previously demonstrated that yeast strains with an overall increase in dNTP pools have much lower levels of rNMPs in their mtDNA than wt yeast^23^, and we hypothesized that the same might be true in mice that lack the SAMHD1 dNTPase^35^, which results in an increase in all dNTPs^28,29^. By keeping dNTP levels low when they are not required for nDNA replication, SAMHD1 protects cells from viral infections^36,37^. Furthermore, mutations in *SAMHD1* are associated with cancer development^38^ and the congenital inflammatory disease Aicardi-Goutières syndrome^39^. As predicted, *SAMHD1*^*−/−*^ mice had far fewer rNMPs in their mtDNA in all tissues examined compared to wt animals. This reduced rNMP content, however, did not improve mtDNA stability in terms of mtDNA deletions, ssDNA breaks, or copy number in aged *SAMHD1*^*−/−*^ animals. Thus, we conclude that the presence of rNMPs in mtDNA cannot account for the mtDNA deletions previously observed in aged animals^3,4^. Furthermore, the lifespan of *SAMHD1*^*−/−*^ animals was similar to their wt littermates, indicating that the “normal” level of rNMPs in wt mtDNA is well tolerated and that the reduction in rNMP content observed in *SAMHD1*^*−/−*^ animals has no significant positive or negative impact. In the future, it will be of great interest to address whether our findings on the mtDNA rNMP content of *SAMHD1*^*−/−*^ animals also hold true for humans, especially in light of the fact that *SAMHD1*^*−/−*^ mice do not exhibit symptoms associated with *SAMHD1* deficiency in humans^27,40^.

Our results also have implications for the understanding of mitochondrial nucleotide metabolism. Mitochondrial dNTP pools are maintained via two routes^41^ – *i)* by the intra-mitochondrial phosphorylation of deoxynucleosides (dNs) that are imported via the equilibrative nucleoside transporter (ENT1) or that result from the degradation of intra-mitochondrial dNTPs (the mitochondrial salvage pathway) and *ii)* via the import of deoxynucleotides (dNMPs, dNDPs, and/or dNTPs) produced in the cytosol. Our finding of reduced rNMPs in the mtDNA of the skeletal muscle, liver, and heart of *SAMHD1*^*−/−*^ mice implies that the import of deoxynucleotides from the cytosol is a major contributor to the size of the mitochondrial dNTP pools in these tissues. Our reasoning is delineated below.

SAMHD1 is a dNTPase that hydrolyses dNTPs to dNs and acts to restrict the size of the cellular dNTP pool outside of S phase^35^. Its deletion therefore results in increased dNTP pools^27–29^. SAMHD1 is mainly localized to the nucleus, with lower amounts found in the cytosol of some cell types, but it is absent in the mitochondria^39,42,43^. Because the mtDNA rNMP content is drastically decreased in *SAMHD1*^*−/−*^ animals, our results demonstrate that changes in the nuclear and/or cytosolic rNTP/dNTP ratios are transmitted into the mitochondria. Because cellular rNTP levels in general show little fluctuation and because SAMHD1 is known to act on dNTPs, it can be assumed that the decreased rNTP/dNTP ratios in *SAMHD1*^*−/−*^ mitochondria are due to increased dNTP – rather than decreased rNTP – levels. In theory, the elevation in mitochondrial dNTPs could be achieved either via intra-mitochondrial phosphorylation of an increased amount of imported dNs (route *i* above) or via the import of an increased amount of deoxynucleotides (route *ii* above). However, because deletion of SAMHD1 leads to elevated dNTPs, rather than elevated dNs, the increased mitochondrial dNTPs in *SAMHD1*^*−/−*^ tissues must be due to the increased cytosolic levels of dNTPs compared to wt animals and can only be observed because import of deoxynucleotides is a major determinant of the size of the mitochondrial dNTP pool. These results are in line with the reported exchange between cytosolic and mitochondrial dNTP pools in cultured cells^44–46^. Our findings therefore demonstrate that also in animal tissues deoxynucleotide import is a major contributor to the size of the mitochondrial dNTP pool.

## Methods

### Animal handling and isolation of embryos and tissues

All mice were maintained at the animal facility at Umeå University under pathogen-free conditions. Mice were housed in an environment with a 12-hour dark/light cycle and *ad libitum* access to food and water. Animal handling and experimental procedure were approved by the ethics committee at Umeå University and complied with the rules and regulations of the Swedish Animal Welfare Agency and with the European Communities Council Directive of 22 September 2010 (2010/63/EU). All efforts were made to minimize animal suffering and to reduce the number of animals used. Homozygous *SAMHD1*^*−/−*^ mice in the C57BL/6 background^27^ were kindly provided by Jan Rehwinkel (University of Oxford, Oxford) and mated with wt C57BL/6 mice. The lifespan study was conducted on 18 wt and 18 *SAMHD1*^*−/−*^ mice. Mice were monitored throughout their lives and humanely sacrificed once they reached a moribund state. The survival of wt vs. knockout animals was compared using the Mantel-Cox test in GraphPad Prism.

For collection of embryos at embryonic day 13.5, the dams were euthanized by cervical dislocation. The uteri were dissected on a petri plate containing ice-cold PBS. The tails of the embryos were isolated for genotyping, and the embryos were snap-frozen in liquid nitrogen and stored at −80°C. For tissues, mice with genotypes of interest were euthanized by CO_2_ inhalation at the following ages: 15 days (“pup”), 13–16 weeks (“adult”), 1 year (“old adult”), or 23–29 months (“aged”). The spleens, livers, hearts, brains, and hind leg muscles (gastrocnemius [GAS] and tibialis anterior [TA]) were placed in Eppendorf tubes, quickly frozen in liquid nitrogen, and kept at −80°C. As far as possible, samples from an equal number of male and female animals were used in each experiment.

### DNA isolation, treatments, agarose gel electrophoresis, and Southern blotting

Individual mouse tissues or whole embryos were incubated overnight with proteinase K (#P6556, Sigma), and total DNA (i.e. nDNA and mtDNA) was isolated according to standard protocols^47^. Where indicated, DNA preparations were digested overnight with the SacI restriction endonuclease followed by treatment with 20 μg RNase A (Thermo Scientific) in the presence of 300 mM NaCl, precipitated in ethanol, and resuspended in TE buffer (10 mM Tris-HCl pH 8.0, 10 mM EDTA). For estimation of rNMP content, DNA was treated with 0.3 M KOH or with RNase HII as previously described^23^.

DNA was separated by electrophoresis on 0.7% agarose gels under either neutral (1× TAE buffer) or denaturing (30 mM NaOH and 1 mM EDTA) conditions at 1.2 V/cm for 16 h at 8°C. The GeneRuler 1 kb DNA ladder (Thermo Scientific) was used as the standard for DNA length. The DNA was transferred to a nylon membrane using standard protocols^47^ and probed sequentially with selected α-dCTP^32^-labelled double-stranded DNA probes, with extensive stripping with boiling 0.1% SDS between probes. The probes used were *COX1* (nt 5,328–6,872) and D-loop (nt 15,652–16,132) for mouse mtDNA and 18S (nt 1,245–1,787 of the 18S rRNA on chromosome 6 [gi 374088232]) for nDNA. In Fig. S1b, end-labeled single-stranded oligos targeting a region of *ND4* (nt 10,430–10,459) in mtDNA were used as strand-specific probes. Blots were exposed to BAS-MS imaging plates (Fujifilm) and scanned on a Typhoon 9400 imager (Amersham Biosciences), and the signal was quantified using ImageJ64 software. The apparent median length of DNA fragments and the rNMP content of mtDNA were determined from the length distributions of untreated and alkali-treated DNA samples as previously described^23^. Because the rNMP content of each individual sample was determined by comparing the median length of untreated vs. alkali-treated aliquots of the same sample, it was not affected by variation in DNA shearing during sample preparation. Data sets for median length and rNMP content were assessed for normal distribution using the Shapiro–Wilk test in GraphPad Prism software, and all tested sample sets passed this test. Statistical comparisons between two groups were performed using Welch’s unequal variances t-test, and each group contained 3–6 animals.

### MtDNA deletion analysis by long-range PCR

The Expand Long Template PCR system (Roche) with forward and reverse primers at nt 2,478–2,512 and nt 1,933–1,906, respectively, was used to amplify a ~15,800 bp fragment of mouse mtDNA from 25 ng of total DNA. The cycling conditions were 92°C 2 min, 30 cycles of (92°C 10 s, 67°C 30 s, 68°C 12 min), 68°C 7 min, and 4°C hold. Products were separated by electrophoresis on a 0.7% agarose gel run in 1× TAE buffer at 55 V in the cold room and imaged on a ChemiDoc Touch instrument (Bio-Rad).

### MtDNA copy number analysis by qPCR

MtDNA copy number was analyzed in duplicate by quantitative real-time PCR using 2 μl of 1/400-diluted SacI-treated total DNA in a 10 μl reaction containing 0.2 μM forward and reverse primers and 1× KAPA SYBR FAST qPCR Master Mix for LightCycler 480 (KAPA Biosystems) in a LightCycler 96 instrument (Roche). Primer pairs targeting cytochrome B (nt 14,682–14,771 of mtDNA^48^) and actin (nt 142,904,143–142,904,053 of chromosome 5 [NC_000071]) were used with the following qPCR program: 95°C 180 s, 40 cycles of (95°C 10 s, 57°C 10 s, 72°C 1 s with signal acquisition), and melting curve (95°C 5 s, 65°C 60 s, heating to 97°C at 0.2°C/s with continuous signal acquisition). C_q_ values determined by the LightCycler 96 software (Roche) were used to calculate the copy number of mtDNA relative to nDNA using the Pfaffl method^49^ and plotted with GraphPad Prism. Statistical comparisons between two groups were performed using Welch’s unequal variances t-test. A total of 5–8 mice were analyzed per genotype.

### Mapping of 5′-ends and rNMPs

The 5′-ends in mtDNA and nDNA from mouse liver were mapped using 5′-End-seq by treating 1 μg total DNA with 0.3 M KCl for 2 h at 55°C. Ribonucleotides were mapped by HydEn-seq^30^ by hydrolyzing 1 μg total DNA with 0.3 M KOH for 2 h at 55°C. Afterwards, ethanol-precipitated DNA fragments were treated for 3 min at 85°C, phosphorylated with 10 U of phosphatase-free T4 polynucleotide kinase (New England BioLabs) for 30 min at 37°C, heat inactivated for 20 min at 65°C, and purified with HighPrep PCR beads (MagBio). Phosphorylated products were treated for 3 min at 85°C, ligated to oligo ARC140^30^ overnight at room temperature with 10 U of T4 RNA ligase, 25% PEG 8000, and 1 mM CoCl_3_(NH_3_)_6_, and purified with HighPrep PCR beads (Mag-Bio). Ligated products were incubated for 3 min at 85°C. The ARC76±ARC77 adaptor was annealed to the fragments for 5 min at room temperature. The second strand was synthesized with 4 U of T7 DNA polymerase (New England BioLabs) and purified with HighPrep PCR beads (MagBio). Libraries were PCR-amplified with KAPA HiFi Hotstart ReadyMix (KAPA Biosystems), purified, quantified with a Qubit fluorometric instrument (Thermo Fisher Scientific), and 75-base paired-end sequenced on an Illumina NextSeq500 instrument to locate the 5′-ends.

### Sequence trimming, filtering, and alignment

All reads were trimmed for quality and adaptor sequence with cutadapt 1.12 (-m 15 -q 10 – match-read-wildcards). Pairs with one or both reads shorter than 15 nt were discarded. Mate 1 of the remaining pairs was aligned to an index containing the sequence of all oligos used in the preparation of these libraries with Bowtie 1.2 (-m1 -v2), and all pairs with successful alignments were discarded. Pairs passing this filter were subsequently aligned to the mm10 *M. musculus* reference genome (-m1 -v2 –X2000). Single-end alignments were then performed with mate 1 of all unaligned pairs (-m1 -v2). Using the –m1 setting causes Bowtie to discard all reads that align to multiple places in the genome, including nuclear mitochondrial DNA segments (NUMTs). To calculate the base identity of 5′-ends or rNMPs in mtDNA and nDNA, the counts of 5′-ends of all paired-end and single-end alignments were determined for all samples and shifted one base upstream to the location of the free 5′-end or hydrolyzed rNMP. The read data were normalized by dividing the reads for each individual rNMP by the strand-specific genome content of its dNMP counterpart. Subsequently, the strand-specific relative proportions of the four rNMPs or 5′-ends were calculated as percentages. Finally, the percentages for the two DNA strands were averaged to generate the numbers representing the genome as a whole.

### Nucleotide pool measurement

For nucleotide pool measurements, mouse embryos and spleens and skeletal muscle from adult mice (aged 13 weeks) were isolated as described above, rapidly placed in 700 μl ice-cold 12% (w/v) TCA and 15 mM MgCl_2_, frozen in liquid nitrogen, and stored at −80°C. Nucleotide extraction was performed as described previously^28^, and sample cleanup over solid-phase extraction columns and high-performance liquid chromatography was carried out as described previously^26,50^.

## Acknowledgements

This project was supported by grants from the Swedish Research Council (to A.C. and to A.R.C.), the Swedish Cancer Society (to A.C.), the Swedish Foundation for Strategic Research (to A.R.C.) and the Lars Hierta Memorial Foundation (to P.H.W.). P.H.W. was supported by grants from the Swedish Cancer Society and the Swedish Society for Medical Research. G.C.D. is supported by the Kempe Foundations. We thank Jan Rehwinkel for the *SAMHD1*^*−/−*^ mice, and acknowledge Dr. Carol Featherstone of Plume Scientific Communication Services for professional editing of the manuscript.

## Author contributions

P.H.W., P.T., and A.C. designed the study; P.H.W., P.T., S.S., K.K., C.N., A.L.F., D.W., and A.K.N. carried out the experiments; P.H.W., L.J.T, S.S., M.K.M.E., A.R.C., and A.C. analyzed the data; P.H.W. and A.C. wrote the paper; and all authors read and edited the paper.

## Competing Interests

The authors declare no competing interests.

## Materials & Correspondence

Correspondence and material requests should be addressed to (P.H.W.) or (A.C.).

## Notes

### Competing Interest Statement

The authors have declared no competing interest.

### Summary of Updates

Figure 6 and the discussion pertaining to mtDNA nicks in the previous version has been removed, and should be considered only preliminary.

## References

1. Gorman, G. S. et al. Mitochondrial diseases. Nat Rev Dis Primers 2, 16080–22 (2016).

2. Cortopassi, G. A. & Arnheim, N. Detection of a specific mitochondrial DNA deletion in tissues of older humans. Nucleic Acids Res 18, 6927–6933 (1990).

3. Pikó, L., Hougham, A. J. & Bulpitt, K. J. Studies of sequence heterogeneity of mitochondrial DNA from rat and mouse tissues: evidence for an increased frequency of deletions/additions with aging. Mech. Ageing Dev. 43, 279–293 (1988).

4. Tanhauser, S. M. & Laipis, P. J. Multiple deletions are detectable in mitochondrial DNA of aging mice. J. Biol. Chem. 270, 24769–24775 (1995).

5. Kennedy, S. R., Salk, J. J., Schmitt, M. W. & Loeb, L. A. Ultra-Sensitive Sequencing Reveals an Age-Related Increase in Somatic Mitochondrial Mutations That Are Inconsistent with Oxidative Damage. PLoS Genet 9, e1003794–10 (2013).

6. Pinto, M. & Moraes, C. T. Mechanisms linking mtDNA damage and aging. Free Radic. Biol. Med. 85, 250–258 (2015).

7. Nick McElhinny, S. A. et al. Abundant ribonucleotide incorporation into DNA by yeast replicative polymerases. Proc Natl Acad Sci USA 107, 4949–4954 (2010).

8. Nick McElhinny, S. A. et al. Genome instability due to ribonucleotide incorporation into DNA. Nat. Chem. Biol. 6, 774–781 (2010).

9. Li, Y. & Breaker, R. R. Kinetics of RNA degradation by specific base catalysis of transesterification involving the 2γ-hydroxyl group. Journal of the American Chemical Society 121, 5364–5372 (1999).

10. Jaishree, T. N., van der Marel, G. A., van Boom, J. H. & Wang, A. H. Structural influence of RNA incorporation in DNA: quantitative nuclear magnetic resonance refinement of d(CG)r(CG)d(CG) and d(CG)r(C)d(TAGCG). Biochemistry 32, 4903–4911 (1993).

11. Chiu, H.-C. et al. RNA intrusions change DNA elastic properties and structure. Nanoscale 6, 10009–10017 (2014).

12. DeRose, E. F., Perera, L., Murray, M. S., Kunkel, T. A. & London, R. E. Solution Structure of the Dickerson DNA Dodecamer Containing a Single Ribonucleotide. Biochemistry 51, 2407–2416 (2012).

13. Sparks, J. L. et al. RNase H2-initiated ribonucleotide excision repair. Mol Cell 47, 980–986 (2012).

14. Reijns, M. A. M. et al. Enzymatic removal of ribonucleotides from DNA is essential for mammalian genome integrity and development. Cell 149, 1008–1022 (2012).

15. Hiller, B. et al. Mammalian RNase H2 removes ribonucleotides from DNA to maintain genome integrity. Journal of Experimental Medicine 209, 1419–1426 (2012).

16. Lujan, S. A., Williams, J. S., Clausen, A. R., Clark, A. B. & Kunkel, T. A. Ribonucleotides are signals for mismatch repair of leading-strand replication errors. Mol Cell 50, 437–443 (2013).

17. Ghodgaonkar, M. M. et al. Ribonucleotides misincorporated into DNA act as strand-discrimination signals in eukaryotic mismatch repair. Mol Cell 50, 323–332 (2013).

18. Pryor, J. M. et al. Ribonucleotide incorporation enables repair of chromosome breaks by nonhomologous end joining. Science 361, 1126–1129 (2018).

19. Wanrooij, P. H. & Chabes, A. Ribonucleotides in mitochondrial DNA. FEBS Lett 121, 5364 (2019).

20. Forslund, J. M. E., Pfeiffer, A., Stojkovič, G., Wanrooij, P. H. & Wanrooij, S. The presence of rNTPs decreases the speed of mitochondrial DNA replication. PLoS Genet 14, e1007315 (2018).

21. Kasiviswanathan, R. & Copeland, W. C. Ribonucleotide Discrimination and Reverse Transcription by the Human Mitochondrial DNA Polymerase. Journal of Biological Chemistry 286, 31490–31500 (2011).

22. Göksenin, A. Y. et al. Human DNA polymerase ε is able to efficiently extend from multiple consecutive ribonucleotides. Journal of Biological Chemistry 287, 42675–42684 (2012).

23. Wanrooij, P. H. et al. Ribonucleotides incorporated by the yeast mitochondrial DNA polymerase are not repaired. Proc. Natl. Acad. Sci. U.S.A. 114, 12466–12471 (2017).

24. Berglund, A.-K. et al. Nucleotide pools dictate the identity and frequency of ribonucleotide incorporation in mitochondrial DNA. PLoS Genet 13, e1006628 (2017).

25. Moss, C. F. et al. Aberrant ribonucleotide incorporation and multiple deletions in mitochondrial DNA of the murine MPV17 disease model. Nucleic Acids Res 45, 12808–12815 (2017).

26. Kong, Z. et al. Simultaneous determination of ribonucleoside and deoxyribonucleoside triphosphates in biological samples by hydrophilic interaction liquid chromatography coupled with tandem mass spectrometry. Nucleic Acids Res 57, 349–8 (2018).

27. Rehwinkel, J. et al. SAMHD1-dependent retroviral control and escape in mice. The EMBO Journal 32, 2454–2462 (2013).

28. Rentoft, M. et al. Heterozygous colon cancer-associated mutations of SAMHD1 have functional significance. Proc Natl Acad Sci USA 113, 4723–4728 (2016).

29. Behrendt, R. et al. Mouse SAMHD1 has antiretroviral activity and suppresses a spontaneous cell-intrinsic antiviral response. Cell Rep 4, 689–696 (2013).

30. Clausen, A. R. et al. Tracking replication enzymology in vivo by genome-wide mapping of ribonucleotide incorporation. Nat Struct Mol Biol 22, 185–191 (2015).

31. Lebedeva, M. A. & Shadel, G. S. Cell cycle- and ribonucleotide reductase-driven changes in mtDNA copy number influence mtDNA Inheritance without compromising mitochondrial gene expression. Cell Cycle 6, 2048–2057 (2007).

32. Grossman, L. I., Watson, R. & Vinograd, J. The presence of ribonucleotides in mature closed-circular mitochondrial DNA. Proc. Natl. Acad. Sci. U.S.A. 70, 3339–3343 (1973).

33. Wong-Staal, F., Mendelsohn, J. & Goulian, M. Ribonucleotides in closed circular mitochondrial DNA from HeLa cells. Biochem Biophys Res Commun 53, 140–148 (1973).

34. Miyaki, M., Koide, K. & Ono, T. RNase and alkali sensitivity of closed circular mitochondrial DNA of rat ascites hepatoma cells. Biochem Biophys Res Commun 50, 252–258 (1973).

35. Franzolin, E. et al. The deoxynucleotide triphosphohydrolase SAMHD1 is a major regulator of DNA precursor pools in mammalian cells. Proc Natl Acad Sci USA 110, 14272–14277 (2013).

36. Laguette, N. et al. SAMHD1 is the dendritic- and myeloid-cell-specific HIV-1 restriction factor counteracted by Vpx. Nature 474, 654–657 (2011).

37. Hrecka, K. et al. Vpx relieves inhibition of HIV-1 infection of macrophages mediated by the SAMHD1 protein. Nature 474, 658–661 (2011).

38. Clifford, R. et al. SAMHD1 is mutated recurrently in chronic lymphocytic leukemia and is involved in response to DNA damage. Blood 123, 1021–1031 (2014).

39. Rice, G. I. et al. Mutations involved in Aicardi-Goutières syndrome implicate SAMHD1 as regulator of the innate immune response. Nat Genet 41, 829–832 (2009).

40. Roesch, F. & Schwartz, O. The SAMHD1 knockout mouse model: in vivo veritas? The EMBO Journal 32, 2427–2429 (2013).

41. Rampazzo, C. et al. Regulation by degradation, a cellular defense against deoxyribonucleotide pool imbalances. Mutat. Res. 703, 2–10 (2010).

42. Baldauf, H.-M. et al. SAMHD1 restricts HIV-1 infection in resting CD4(+) T cells. Nat. Med. 18, 1682–1687 (2012).

43. Brandariz-Nuñez, A. et al. Role of SAMHD1 nuclear localization in restriction of HIV-1 and SIVmac. Retrovirology 9, 49–12 (2012).

44. Pontarin, G., Gallinaro, L., Ferraro, P., Reichard, P. & Bianchi, V. Origins of mitochondrial thymidine triphosphate: dynamic relations to cytosolic pools. Proc. Natl. Acad. Sci. U.S.A. 100, 12159–12164 (2003).

45. Ferraro, P. et al. Mitochondrial deoxynucleotide pools in quiescent fibroblasts: a possible model for mitochondrial neurogastrointestinal encephalomyopathy (MNGIE). J. Biol. Chem. 280, 24472–24480 (2005).

46. Leanza, L., Ferraro, P., Reichard, P. & Bianchi, V. Metabolic interrelations within guanine deoxynucleotide pools for mitochondrial and nuclear DNA maintenance. J. Biol. Chem. 283, 16437–16445 (2008).

47. Sambrook, J. & Russell, D. W. Molecular Cloning. (CSHL Press, 2001).

48. Ahola-Erkkilä, S. et al. Ketogenic diet slows down mitochondrial myopathy progression in mice. Human molecular genetics 19, 1974–1984 (2010).

49. Pfaffl, M. W. A new mathematical model for relative quantification in real-time RT-PCR. Nucleic Acids Research 29, e45 (2001).

50. Jia, S., Marjavaara, L., Buckland, R., Sharma, S. & Chabes, A. Determination of deoxyribonucleoside triphosphate concentrations in yeast cells by strong anion-exchange high-performance liquid chromatography coupled with ultraviolet detection. Methods Mol Biol 1300, 113–121 (2015).

