## Supplemental information for "The physiological level of rNMPs present in mtDNA does not compromise its stability"

**Supplementary figure legends**

**Figure S1. Related to Figure 1. a)** The blot shown in Fig. 1b was stripped and re-probed with a dsDNA probe for the D-loop region. The 7S DNA is indicated (7S). **b)** The blot shown in Fig. 1b was stripped and re-probed with strand-specific probes for the *ND4* gene in mtDNA: probes H4 (left panel) and L4 (right panel) detect the L- and H-strands, respectively. **c)** The blot shown in Fig. 1b was stripped and re-probed with a dsDNA probe for the 18S rDNA region in nDNA. The size of bands in the DNA ladder are indicated in kb.

**Figure S2. Related to Figure 2. a)** The Southern blot shown in Fig. 2a was stripped and re-probed with a dsDNA probe against the 18S rDNA region of nDNA. **b)** Untreated DNA and KOH-treated DNA from the hearts of pups or adult mice was separated by electrophoresis on a denaturing gel and mtDNA was visualized using a *COX1* probe. Each pair of untreated and KOH-treated sample was run on the same gel. Dotted lines represent the median. **c)** The Southern blot shown in Fig. S2b was stripped and re-probed with a dsDNA probe for the 18S rDNA region in nDNA. The size of bands in the DNA ladder are indicated in kb.

**Figure S3. Related to Figure 3. a)** The Southern blot shown in Fig. 3a was stripped and re-probed with a dsDNA probe against the 18S rDNA region in nDNA. **b)** The Southern blot shown in Fig. 3c was stripped and re-probed with a dsDNA probe against the 18S rDNA region in nDNA. **c)** Untreated and KOH-treated DNA from the heart muscle of old adult wt, *SAMHD1*$^{+/-}$ (het) and *SAMHD1*$^{-/-}$ (ko) mice was separated by electrophoresis on a denaturing gel and mtDNA was visualized by Southern blotting using a *COX1* probe. Each lane corresponds to an individual mouse. **d)** The length difference between untreated and KOH-treated mtDNA shown in Fig. S3c was used to compute the number of rNMPs per single-strand of mtDNA. The p-values of statistically significant differences between the wt and other groups are indicated (Welch's t-test; n=3). The horizontal lines indicate the median for each group. **e)** The Southern blot shown in Fig. S3c was stripped and re-probed with a dsDNA probe for the 18S rDNA region in nDNA. The size of bands in the DNA ladder are indicated in kb.

**Figure S4. Related to Figure 4. a)** 5′-end sequencing was used to identify the proportion of each base (A, T, G, C) at the 5′ ends of mtDNA from the livers of adult wt and *SAMHD1*$^{-/-}$ (ko) mice. The same DNA samples as those in Fig. 4a were analysed, but samples were treated with KCl instead of KOH. Error bars represent the standard deviation. **b)** The proportions of rNMPs in the reverse (R) and forward (F) strands of nDNA in wt and *SAMHD1*$^{-/-}$ (ko) mice after normalization to the base content of nDNA. Error bars represent the standard deviation. **c)** The proportion (%) of each rNMP in each nDNA strand (R, F) in wt and ko mice. Error bars represent standard deviation.

**Figure S5. Related to Figure 5. a)** MtDNA copy number was determined by qPCR of DNA from the livers of 5–8 wt (filled dots) and *SAMHD1*$^{-/-}$ (ko, open dots) mice at different ages; copy number values are expressed as % of the adult wt value. The mean is indicated by a line. The p-values were calculated by using Welch's t-test: ns, non-significant. **b)** The blot shown in Fig. 5d was stripped and re-probed with a dsDNA probe for the 18S rDNA region in nDNA. The size of the bands in the DNA ladder are indicated in kb.

### Supplemental Figure 1; related to Fig. 1

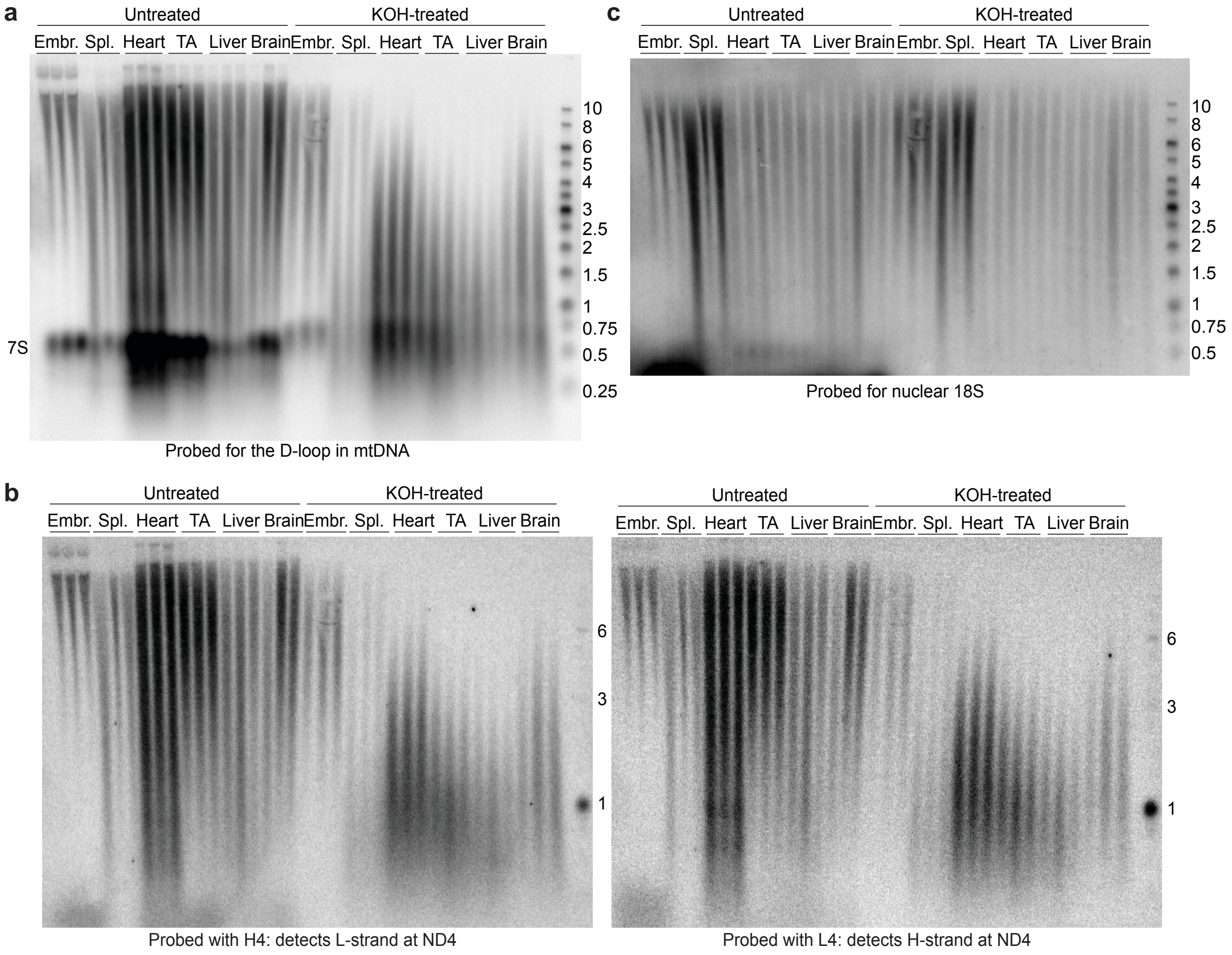

**Supplemental Figure 2; related to Fig. 2**

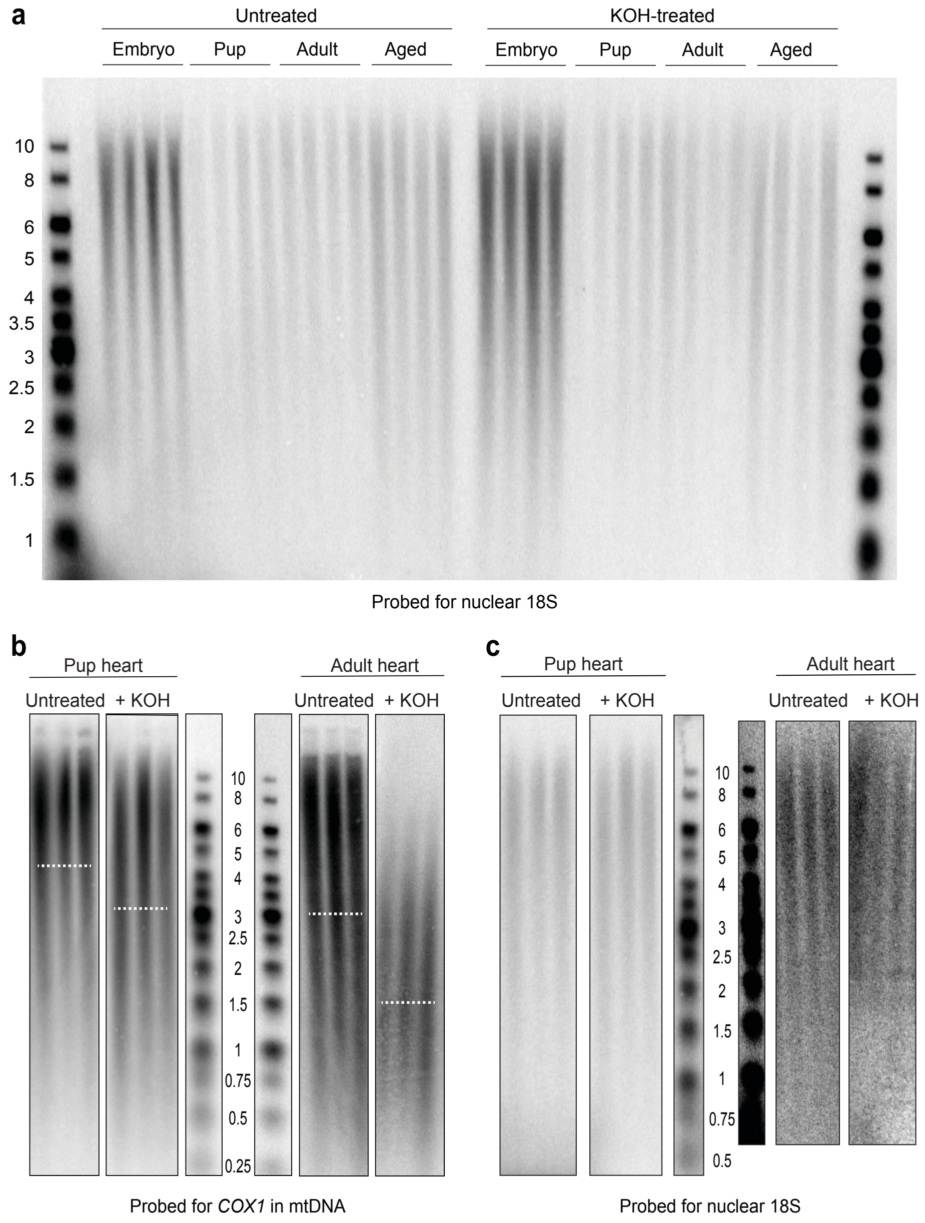

### Supplemental Figure 3; related to Fig. 3

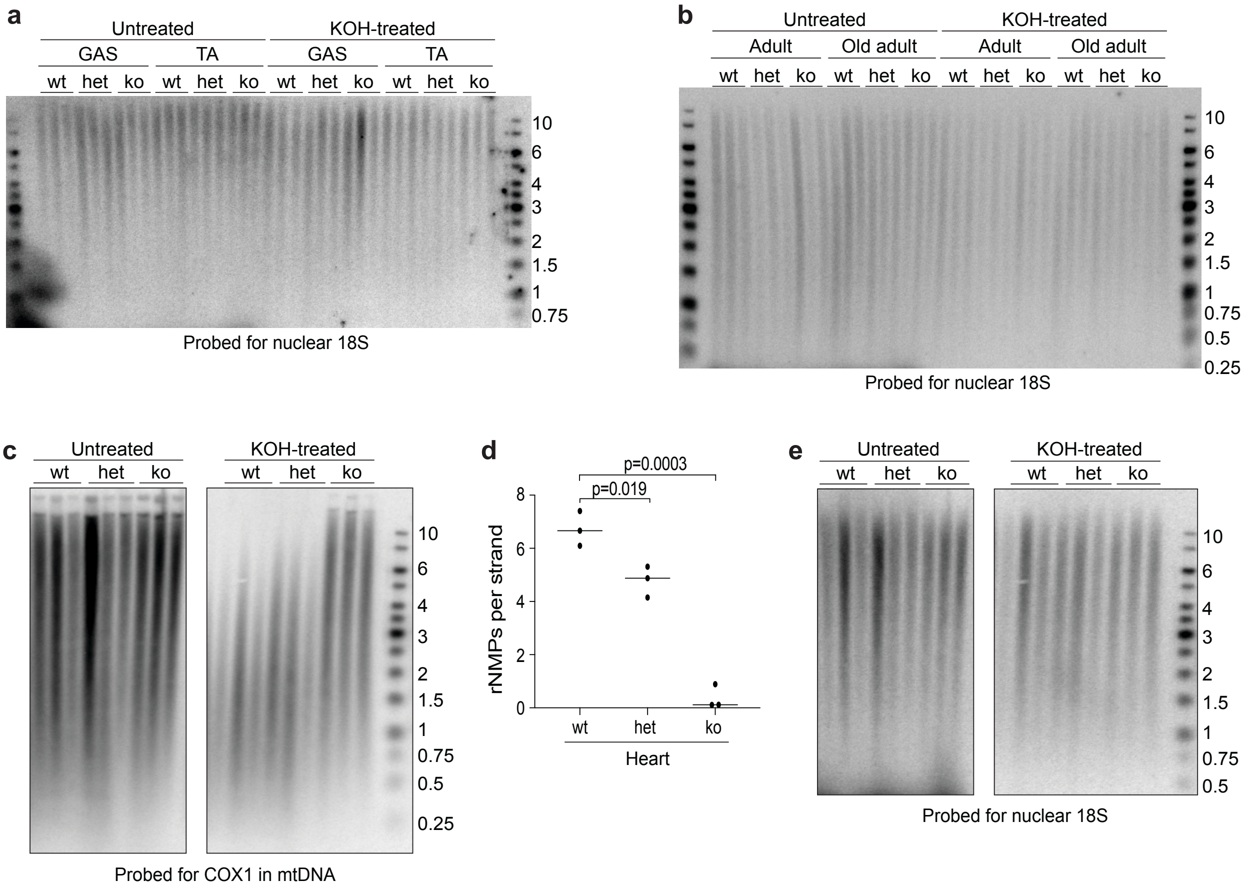

### Supplemental Figure 4; related to Fig. 4

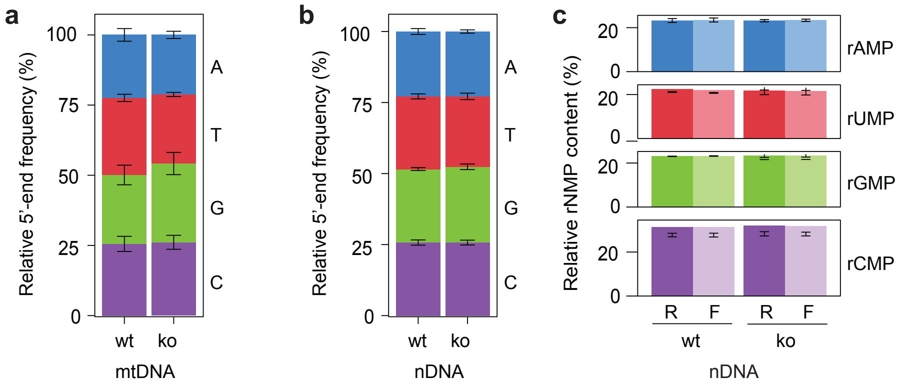

### Supplemental Figure 5; related to Fig. 5

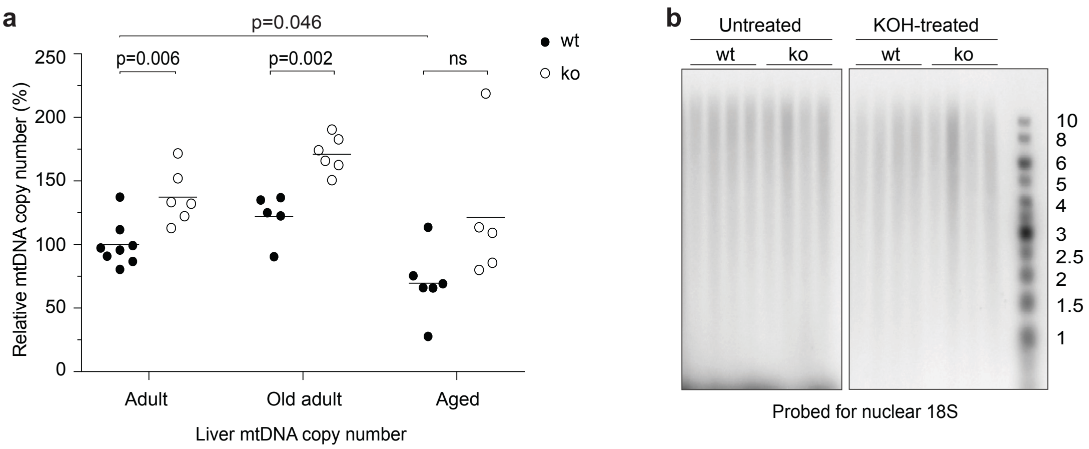
